## supplemental figures for "Cardiovirus leader proteins retarget RSK kinases toward alternative substrates to perturb nucleocytoplasmic traffic"

Supplemetary figure legends

**Figure S1. BioID-RSK and BioID-L fusion proteins keep their normal activities.**

(A) L protein's activities are conserved in HeLa BioID-RSK cells. HeLa BioID-RSK cells were infected with $L^{WT}$ and $L^{M60V}$ viruses for 16h. Western blot showing BioID-RSK activation by the $L^{WT}$ and $L^{M60V}$ proteins (p-RSK at S380) and PKR inhibition by LWT but not by $L^{M60V}$ (p-PKR at T446 is a marker of PKR activation). (B) BioID-L fusion proteins maintain their corresponding activities. Immunoblots show the detection of activated RSK (p-S380), activated PKR (p-T446) and Nup98 hyperphosphorylation (migration shift) in HeLa cells infected for 16h with BioID-$L^{WT}$, BioID-$L^{M60V}$, BioID-$L^{F48A}$ replicons (2 lanes each) and, as a control, with $L^{WT}$ and $L^{M60V}$ viruses (1 lane each). Viral polymerase 3D and ß-actin were detected as infection and loading controls respectively.

**Figure S2. Analog-sensitive RSKs (As1-RSK and As2-RSK) keep their kinase activity and thiophosphorylate substrates in vitro.**

(A) Analog-sensitive RSKs (As1 and As2-RSKs) recue L protein activities. HeLa RSK-TKO cells transduced with an empty vector (-) or with lentiviral vectors expressing WT-RSK, As1-RSK or As2-RSK, were infected (MOI 2.5) for 15h with $L^{WT}$ or $L^{M60V}$ viruses. Western blots show the detection of HA-(RSK), P-PKR (T446), PKR, NUP98, 3D viral polymerase as a control of infection and ß-actin as a loading control. RSKs activation (p-RSK S380) PKR inhibition (inhibition of pPKR T446) and NUP98 hyperphosphorylation (shift upwards) in As-RSK expressing cells paralleled those observed in WT-RSK expressing cells. (B) GST-S6 (thio)-phosphorylation by As1, As2 and WT RSKs in an in vitro kinase assay. 293T cells were transfected with plasmids coding for WT-RSK, as1-RSK or as2-RSK. 6 hours post-transfection, cells were treated with phorbol myristate acetate (PMA) to activate RSKs. 18 hours later, RSKs were immunoprecipitated with an anti-HA antibody. An in vitro kinase assay was performed with the immunoprecipitated RSKs, GST-S6 (recombinant substrate) and either ATP or N6-Bn-ATP-$\gamma$-S or N6-PhEt-ATP-$\gamma$-S. Reaction proceeded for 30min at 30°C before alkylation by PNBM for 2h at room temperature. Reaction was stopped by addition of sample buffer. Samples were analyzed by western blot with antibodies against HA-(RSK), RxxS*/T* (antibody against phosphorylated RSK substrates; here: phospho-GST-S6), and against thiophosphate ester. WT RSK was able to use ATP and phosphorylate GST-S6 (RxxS*/T*) but was not able to use the ATP analogs and thiophosphorylate (Thiophosphate ester). As1 and As2-RSK were able to use ATP and phosphorylate GST-S6 (RxxS*/T*) but were also able to use both ATP analogs, with a preference for N6-Bn-ATP-$\gamma$-S (Thiophosphate ester). Dashed lines between lanes indicate deletion of irrelevant sections from the same membrane. (C) Thiophosphorylation of NUP214 by As2-RSK. Immunoblots showing NUP214 in the thiophosphate ester IP fraction when $L^{WT}$ is present. HeLa cells expressing As2-RSK or WT-RSK were infected with TMEV for 8h (MOI 5). Cells were then permeabilized with digitonin, and N6-Bn-ATP-$\gamma$-S was added for 1 hour. Cells were then lysed and thiophosphate-ester containing proteins were immunoprecipitated.

Fig. S1

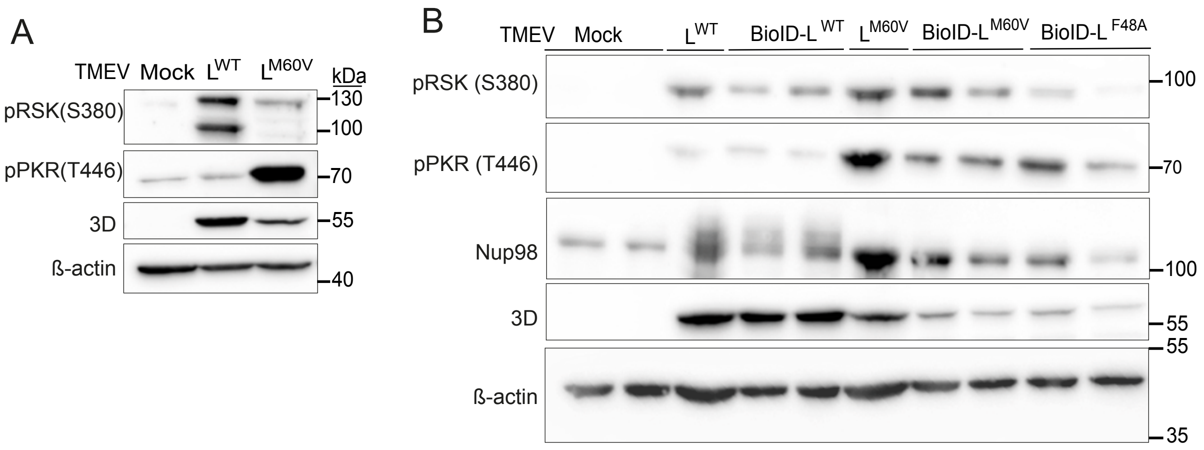

A
TMEV Mock L WT L M60V
kDa
pRSK(S380)
130
100
pPKR(T446)
70
3D
55
ß-actin
40
B
TMEV Mock L WT BioID-L WT L M60V BioID-L M60V BioID-L F48A
pRSK (S380)
100
pPKR (T446)
70
Nup98
100
3D
55
55
ß-actin
35

Fig. S2

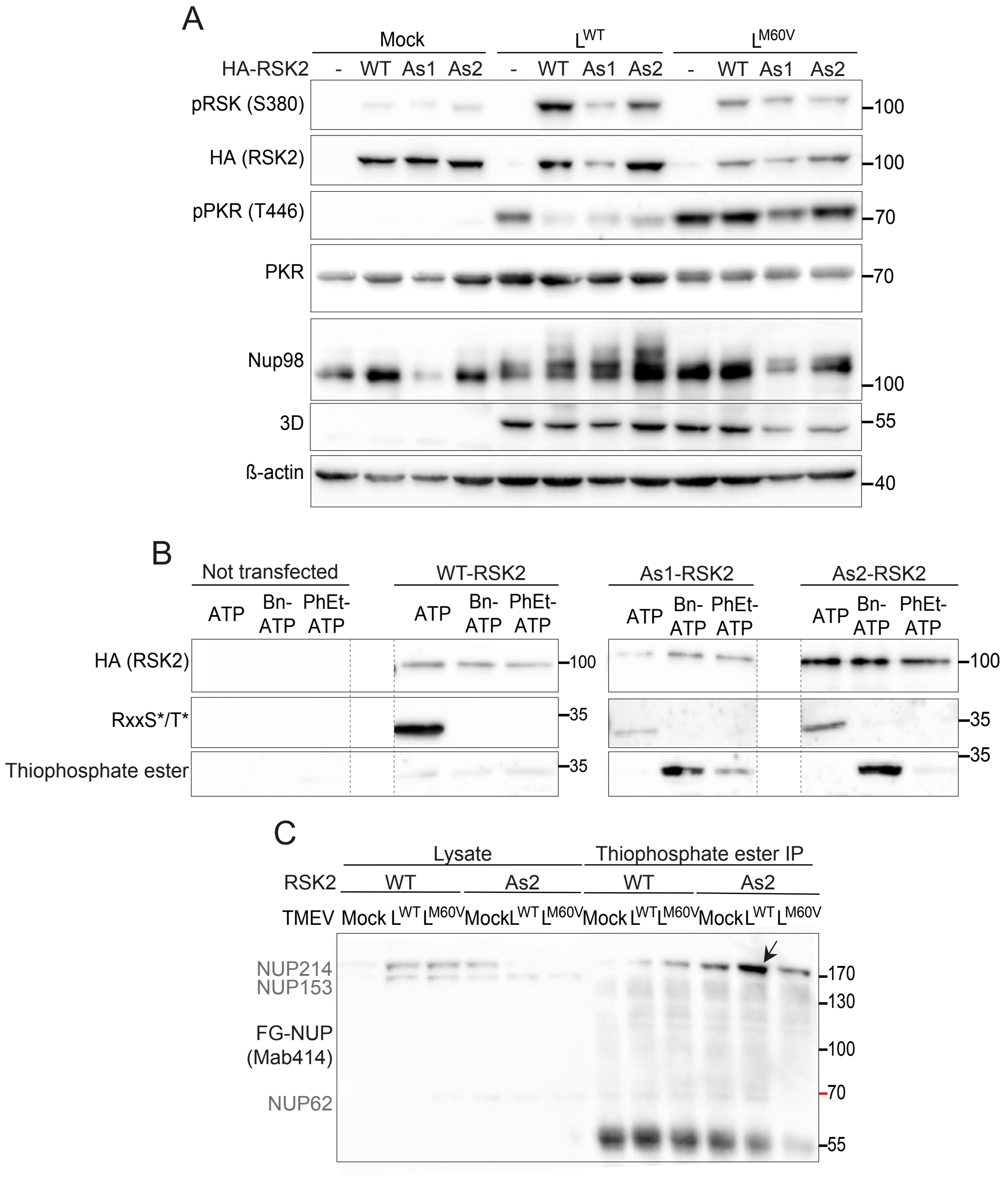
A
Mock
L^WT
L^M60V
HA-RSK2
-
WT
As1
As2
pRSK (S380)
HA (RSK2)
pPKR (T446)
PKR
Nup98
3D
ß-actin
100
70
55
40
B
Not transfected
WT-RSK2
As1-RSK2
As2-RSK2
ATP
Bn-ATP
PhEt-ATP
HA (RSK2)
RxxS*/T*
Thiophosphate ester
100
35
C
Lysate
Thiophosphate ester IP
RSK2
WT
As2
TMEV
Mock
L^WT
L^M60V
NUP214
NUP153
FG-NUP (Mab414)
NUP62
170
130
100
70
55
